# Transgenerational dispersal plasticity and its fitness consequences are under genetic control

**DOI:** 10.1101/791210

**Authors:** Hugo Cayuela, Staffan Jacob, Nicolas Schtickzelle, Rik Verdonck, Hervé Philippe, Martin Laporte, Michèle Huet, Louis Bernatchez, Delphine Legrand

## Abstract

Phenotypic plasticity, the ability of one genotype to produce different phenotypes in different environments, plays a central role in species’ response to environmental changes. Transgenerational plasticity (TGP) allows the transmission of this environmentally-induced phenotypic variation across generations, and can influence adaptation. To date, the genetic control of TGP, its long-term stability, and its potential costs remain largely unknown, mostly because empirical demonstrations of TGP across many generations in several genetic backgrounds are scarce. Here, we examined how genotype determines the TGP of dispersal, a fundamental process in ecology and evolution. We used an experimental approach involving ~200 clonal generations in a model-species of ciliate to determine if and how TGP influences the expression of dispersal-related traits in several genotypes. Our results show that morphological and movement traits associated with dispersal are plastic, and that these modifications are inherited over at least 35 generations. We also highlight that genotype modulates the fitness costs and benefits associated with plastic dispersal strategies. Our study suggests that genotype-dependent TGP could play a critical role in eco-evolutionary dynamics as dispersal determines gene flow and the long-term persistence of natural populations. More generally, it outlines the tremendous importance that genotype-dependent TGP could have in the ability of organisms to cope with current and future environmental changes.

**Significance:** The genetic control of the transgenerational plasticity is still poorly understood despite its critical role in species responses to environmental changes. We examined how genotype determines transgenerational plasticity of a complex trait (*i.e.*, dispersal) in a model-species of ciliate across ~200 clonal generations. Our results provide evidence that plastic phenotypic variation linked to dispersal is stably inherited over tens of generations and that cell genotype modulates the expression and fitness cost of transgenerational plasticity.

## Introduction

Transgenerational plasticity (TGP) is a central mechanism in the evolution of the living world (Uller 2008, Herman & Sultan 2011). TGP occurs when abiotic (e.g., Galloway & Etterson 2007, Marshall 2008, Heckwolf et al. 2018) and biotic (Dantzer et al. 2013) environmental conditions alter the phenotype of parents and when those changes then affect offspring phenotypic expression. For instance, parents can produce young with phenotypic characteristics that increase their fitness when exposed to similar environmental conditions (*i.e.*, adaptive TGP; e.g., Dantzer et al. 2013). Alternatively, phenotypic modifications induced by TGP may decrease offspring performance via transgenerational costs (*i.e.*, maladaptive TGP; e.g., Marshall 2008). The ability to transmit and express an advantageous phenotype in the next generation(s), or to mitigate the costs of TPG, could depend on the genetic background (Herman & Sultan 2016), similar to phenotypic plasticity in general. Indeed, evolution of reaction norms (slope and curvature) and the mitigation of plastic costs can depend on specific genetic variants (*i.e.*, G × E interactions; Gerken et al. 2015), epigenetic marks under strict or partial genetic control (Kooke et al. 2015) and the regulation of gene expression (Murren et al. 2015). However, with the exception of the predictions from a handful of theoretical models (Greenspoon & Spencer 2018), the role of genetic background in TGP evolution remains poorly understood despite its critical importance for the ability of the living to cope with current global change (Guillaume et al. 2016, Donelson et al. 2017).

Dispersal, the movement of individuals potentially leading to gene flow (Ronce 2007), is a highly relevant candidate for investigating TGP mechanisms. Dispersal is a complex and multidimensional phenotype, which is highly plastic at all its stages (emigration, transience, and emigration; Clobert et al. 2009, Cote et al. 2017) and under partial genetic control (Saastamoinen et al. 2018). Its evolution is determined by the balance between the fitness benefit of moving (for instance, to escape local detrimental conditions for survival or reproduction) and the related costs (Clobert et al. 2009, Bonte et al. 2012). Dispersal is especially constrained by direct (*e.g.*, energy and time) costs incurred during the displacements in the landscape matrix and indirect costs associated with the expression of phenotypic traits facilitating dispersal (Bonte et al. 2012). These associations between dispersal and other traits are called “dispersal syndromes” (Ronce & Clobert 2012) and may result in trade-offs when traits are negatively correlated with fitness components, notably due to gene pleiotropy (Saastamoinen et al. 2018). Studies have suggested that TGP may facilitate the transmission of traits across generations that improve dispersal in a given environmental context (Bitume et al. 2015), while offering the possibility to reverse or explore other phenotypic states if the environment changes again (Saastamoinen et al. 2018). In absence of empirical evidence, one might expect that TGP for dispersal could occur in concert with the transmission of its fitness consequences across generations. In addition, the genetic background of parents could affect the ability to transmit dispersal-related traits and could modulate fitness costs associated to the expression of those traits across generations. However, these hypotheses have not been yet tested due to difficulties in studying TGP across many generations and across different genotypes.

Here, we investigated the genetic control of TGP for dispersal-related traits and the related fitness consequences in the protist *Tetrahymena thermophila*. This species reproduces clonally in standard laboratory conditions (Bell & Stein 2017), with the availability of several genotypes showing different degrees of dispersal plasticity (*e.g.*, Schtickzelle et al. 2009, Pennekamp et al. 2014, Jacob et al. 2016). It thus represents an excellent biological model to study TGP for dispersal. We used a procedure of successive dispersal trials in controlled microcosms to produce two cell lines, dispersing *vs* non-dispersing cells, in four isogenic strains (*i.e*. negligible genetic variation inside a strain) thereafter called “D3, D4, D6 and D9” (**Supplementary material**, **Fig.S1**). To control for genetic variation while testing whether TGP explains experimental patterns, mother cultures were established from the isolation of a single cell for each genotype, and these cultures were then split into five replicates. Experiments were also limited to six weeks with one dispersal trial per week (~200 asexual generations for the entire experiment). This procedure prevents the possibility that pre-existing genetic variation explains the observed phenotypic pattern during experiment and excludes a major role of new genetic variation. Before the first dispersal trial, we verified the degree of genetic control in a set of morphological (cell size and shape) and movement (velocity and linearity) traits related to dispersal in *T. thermophila*, as well as in fitness using cell growth as a proxy (*e.g.*, Orr 2009). Then, during the first dispersal trial, we examined if dispersing and non-dispersing cells differ in their morphological and movement traits, resulting in the existence of a plastic dispersal syndrome (*e.g.*, Ronce & Clobert 2012, Stevens et al. 2013, Legrand et al. 2016). Next, we investigated how the genetic background determines the plastic response of these dispersal-related traits during the six successive dispersal trials (separated by ~35 cell divisions). We especially tested the hypotheses that (1) the dispersal status of ancestors affects the phenotype of descendants across several generations (existence of TGP), and that (2) the strength of immediate and transgenerational plastic response varies with the genetic background. We also examined (3) if the observed TGP was gradual or stable when repeated dispersal trials are experienced by ancestors (e.g., Vastenhouw et al. 2006, Remy 2010, Sentis et al. 2018). Finally, we tested that (4) dispersing cells incur a fitness cost (Bonte et al. 2012) at the first dispersal trial and whether this cost is cumulative through generations and modulated by the genotype.

## Results

### Trait covariation, fitness and dispersal syndrome after the first trial

Before the initial dispersal trial (*tr*0, see **Fig.S1**), we examined the individual covariation among the four tested dispersal-related traits (**models 1 of Table S1**). Here we show only significant associations with an explained variation higher than 1% (based on R^2^), which we consider as potentially biologically relevant. We found that velocity was positively correlated to movement linearity (*R*^2^ = 0.05, *χ*^2^ = 2533.10, *p* < 0.0001) and cell shape (*R*^2^ = 0.16, LR test: X^2^ = 11926.00, *p* < 0.0001; the fastest cells had the most linear movements and the most elongated cells were the fastest). Furthermore, cell shape and movement linearity were positively related (*R*^2^ = 0.02, *χ*^2^ = 1996.40, *p* < 0.0001; the most elongated cells had the most linear movements, see **Table S3** for all relationships). In addition to these four phenotypic traits, we measured cell growth rate estimated from 15 days (~75 generations), a common fitness proxy in *T. thermophila*. Growth rate was negatively correlated to cell shape (*R*^2^ = 0.45, *χ*^2^ = 4.15, *p* = 0.04), but no significant relationship was found with cell size (*R*^2^ = 0.14, *χ*^2^ = 1.21, *p* = 0.27), linearity (*R*^2^ = 0.08, *χ*^2^ = 0.69, *p* = 0.40) and velocity (*R*^2^ = 0.12, *χ*^2^ = 0.98, *p* = 0.32).

We then examined the effect of genotype identity on the four phenotypic traits **(models 2 of Table S1)**. Genotype explained 46% of cell size variation (*χ*^2^ = 90.31, *p* < 0.0001), 26% of cell shape variation (*χ*^2^ = 64.67, *p* < 0.0001), 7% of movement linearity variation (*χ*^2^ = 49.39, *p* < 0.0001), and 5% of velocity variation (*χ*^2^ = 11.47, *p* = 0.009). Furthermore, we showed that genotype identity explained 87% of variation in growth rate (*χ*^2^ = 44.65, *p* < 0.0001). These results indicate strong phenotypic differences between the genetic backgrounds used in our experiments.

Next, we investigated dispersal syndrome by performing an immediate quantification of the association between dispersal and phenotypic traits just after the first dispersal trial (*tr*0; see **models 3 of Table S1**). Dispersing cells were more elongated (*R*^2^ = 0.02, *χ*^2^ = 61.94, *p* < 0.0001) and swam faster (*R*^2^ = 0.03, *χ*^2^ = 77.69, *p* < 0.0001) than non-dispersing cells. By contrast, dispersing and non-dispersing cells did not significantly differ in terms of size (*R*^2^ = 0.001, *χ*^2^ = 2.18, *p* = 0.13) and movement linearity (*R*^2^ = 0.001, *χ*^2^ = 2.14, *p* = 0.16). Fitness differed between the dispersing and non-dispersing cells of each genotype: dispersing cells had lower growth rate than non-dispersing ones (*R*^2^ = 0.04, *χ*^2^ = 15.48, *p* < 0.0001), indicating a dispersal-related fitness cost.

Altogether, our results highlight the existence of trait-trait correlations and a plastic dispersal syndrome. As cell size and movement linearity marginally differed between dispersing and non-dispersing cells, we focused further analyses on cell velocity and shape, the two traits that most contributed to dispersal.

### Effect of ancestor dispersal status and genotype on descendant phenotype across generations

Following each dispersal trial, we first tested for the persistence of trait divergence between dispersing and non-dispersing cells after ~35 asexual generations in common garden conditions in the whole dataset (**models 1.1 of Table S2**; **Fig S1**). The dispersal status of cell ancestors, *i.e.* cells from the dispersing *vs* non-dispersing selected lines, affected descendants’ velocity (*R*^2^ = 0.05, *χ*^2^ = 8.58, *p* = 0.003) and shape (*R*^2^ = 0.01, *χ*^2^ = 8.58, *p* = 0.03). Cells with a dispersing ancestor recurrently had a higher velocity and a more elongated shape than those with a non-dispersing ancestor (Fig.1A and Fig.1B).

**Fig. 1.**
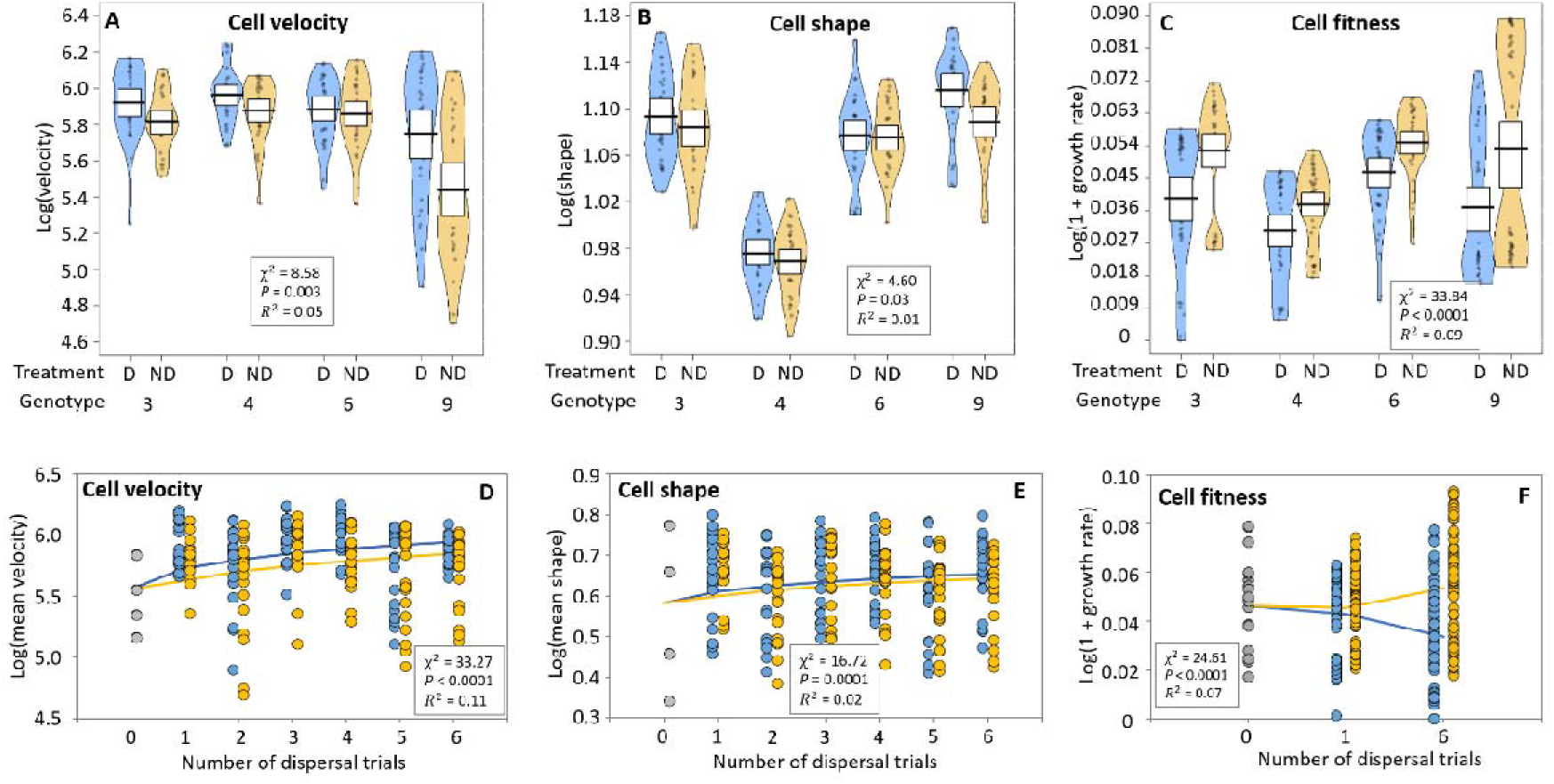
Transgenerational plasticity for dispersal and its fitness cost: effect of ancestor dispersal status (dispersing ancestor in blue and non-dispersing ancestor in yellow) on phenotypic traits (cell shape and velocity) and fitness of descendants ~35 asexual generations after dispersal trials (i.e., just before the next one) in the four studied genotypes (D3, D4, D6, and D9). Mother cultures are represented in grey. (A-B-C) We show relationships where the effect of the ancestor dispersal status on phenotypic traits was significant with a p-value threshold of p = 0.05; non-significant relationships are shown in **Supplementary material, Fig. S2**. We provide marginal of the mixed model and outputs of the likelihood ratio test (and P-value) used to examine the effect of dispersal trial on phenotypic traits. (D-E-F) Effect of the number of dispersal trials experienced by ancestors on cell phenotype. The terms ‘ancestor dispersal status’ and ‘number of trials’ were entered in an additive way in the model (the interaction was not supported by the data). We give marginal of the sum of fixed effects in the mixed model and outputs of the likelihood ratio test used to examine the effect of number of dispersal trials on phenotypic traits.

Second, we examined how the strength of the effect of ancestor dispersal status on phenotypic traits differed among genotypes (**models 1.2 of Table S2** and model outputs presented in **Table S5**). The ancestor dispersal status explained from 0.3% to 13% of velocity variation in D6 and D9 respectively, and from 0.1 and 11% of shape variation in D6 and D9 respectively. Cells with dispersing ancestors had higher dispersal rates than cells with non-dispersing ancestors in D9 (*R*^2^ = 0.06, *χ*^2^ = 6.65, *p* = 0.01); the effect was marginal in D3 (*R*^2^ = 0.04, *χ*^2^ = 3.28, *p* = 0.07) and D6 (*R*^2^ = 0.04, *χ*^2^ = 3.12, *p* = 0.08), and not significant in D4 (*R*^2^ = 0.02, *χ*^2^ = 1.39, *p* = 0.23).

Increasing number of dispersal trials experienced by each experimental line did not cause a gradual change of trait values with time (**models 2 of Table S2)**. The shape and velocity differences between the descendants of dispersing and non-dispersing cells appeared at *tr*0 and did not increase nor decrease over the following trials (from *tr*1 to *tr*6, Fig.1D and Fig.1E). Accordingly, the association between these phenotypic traits and the number of dispersal trials was better described by a logarithmic relationship than a linear relationship (velocity, *χ*^2^ = 35.40, *p* < 0.0001; shape, *χ*^2^ = 18.35, *p* = 0.0001). In addition, the interaction between ‘ancestor dispersal status’ and ‘number of trials’ was not supported by the data for the two phenotypic traits (**Table S4**).

We then tested how stable these transgenerational changes of cell phenotype were by comparing the phenotype of cells measured after each dispersal trial and the phenotype of their descendants after ~ 35 asexual generations in common garden (**model 3 of Table S2**). Cells with a dispersing ancestor had a lower velocity ~35 generations after the trial than immediately after the trial (*R*^2^ = 0.23, *χ*^2^ = 82.09, *p* < 0.0001), indicating that this trait was partially reversible under standard environmental conditions. Yet, the reversibility was not sufficiently strong to eliminate the effect of ancestor dispersal status on descendant phenotype (Fig.1A). By contrast, the shape of descendants was more elongated than that of their ancestor (= 0.09, = 52.19, *p* < 0.0001), suggesting a slight exacerbation of this trait after ~35 generations.

### Effect of genotype and ancestor dispersal status on descendant fitness through time

We examined growth rates of dispersing and non-dispersing lines at three dispersal trials (*tr*0, *tr*1 and *tr*6; see **model 1.1 of Table S2** and **Fig S1**). Pooling these three times and the four genotypes revealed that cells with a dispersing ancestor had a lower growth than those with a non-dispersing ancestor (= 0.09, = 33.84, *p* < 0.0001, Fig 1C), which indicates a transgenerational fitness effect of dispersal trials on descendants. Looking at temporal trends revealed that growth of cells with dispersing ancestors decreased between *tr*0 and *tr*1 and between *tr*1 and *tr*6 while it increased in cells with non-dispersing ancestors (Fig 1F, = 24.61, *p* < 0.0001; **model 2 of Table S2**).

An analysis of the data per genotype showed that the transgenerational fitness cost was modulated by the genetic background (**model 1.2 of Table S2**). Although cells with a dispersing ancestor all experienced a fitness loss, variation explained by the genotype (dispersing vs non-dispersing line) was more important for D3 and D6 (13% and 10% respectively) than for D9 and D4 (6% and 5% respectively) (**Table S6**).

Finally, we found that the fitness consequences of dispersal were weakly reversible as growth rate was similar just after dispersal trials and ~35 generations later for both dispersing (= 0.004, = 2.03, *p* = 0.15) and non-dispersing cells (= 0.001, = 0.45, *p* = 0.50).

## Discussion

During the initial dispersal trial, our results confirmed the existence of a plastic dispersal syndrome in *T. thermophila*. Within the four genotypes, dispersing cells had a more elongated shape and a higher velocity than non-dispersing cells, those two traits generally facilitating dispersal in this species (Fjerdingstad et al. 2007, Pennekamp et al. 2014, Jacob et al. 2016). The difference of velocity and shape between dispersing and non-dispersing cells differed among genotypes. Furthermore, cells with a dispersing phenotype experienced a fitness loss that was modulated by the genotype, corroborating the results of previous studies in *T. thermophila* (Schtickzelle et al. 2009, Jacob et al. 2016). We also confirmed that the described dispersal syndrome and costs result from strong G × E interactions and that intragenerational plasticity is an important driver in dispersal evolution of *T. thermophila* (Pennekamp et al. 2014) and beyond (Saastamoinen et al. 2018). Except for our fitness proxy, the relationships between the dispersal strategies and the correlated traits were weak in our conditions. In our trials, dispersal cues perceived by cells were mainly linked to changes in density and spatial conformation of habitats, two important drivers of dispersal (pipetting of ~100,000 cells from mother cultures kept in a 2mL-well placed in a fresh and empty 1.5mL tube connected by a thin corridor to an empty-of-cell arrival tube). Further experiments in conditions where dispersal might be more beneficial (*e.g.*, temperature or chemical stress, interspecific competition) should inform on the context-dependency of the highlighted dispersal syndrome, especially for the lability and strength of trait correlations (Cote et al. 2017).

Our study demonstrated that plastic phenotypic variation linked to dispersal is stably inherited when cells are exposed to successive dispersal trials separated by ~35 asexual generations (Fig.2). Cells conserved the phenotypic characteristics (shape and velocity) associated with the dispersal status of their ancestors. Our experimental protocol allows us to reasonably assume that the detected phenotypic variation in the descendants results from TGP rather than in genic selection. Indeed, we have eliminated most genetic variation within each replicate at the beginning of the experiment using a single mother cell, which rules out the possibility of selection from standing genetic variation (see further considerations in **Supplement**). We also believe very unlikely that *de novo* mutations have been simultaneously recruited in the four genotypes during the 7-days growth period preceding the first dispersal trial. As a result, the phenotypic changes observed after the first dispersal trial, and maintained at least during ~35 generations, are due to transgenerational plastic mechanisms. Examples of TGP observed for more than a few generations are not frequent and mostly found in other (partially) asexual species (Vastenhouw et al. 2006). Here, we demonstrate that TGP over tens of asexual generations can influence dispersal, an eco-evolutionary force that could act to enhance gene flow. Future research should determine if TGP for dispersal occurs also across sexual generations in this ciliate.

**Fig. 2.**
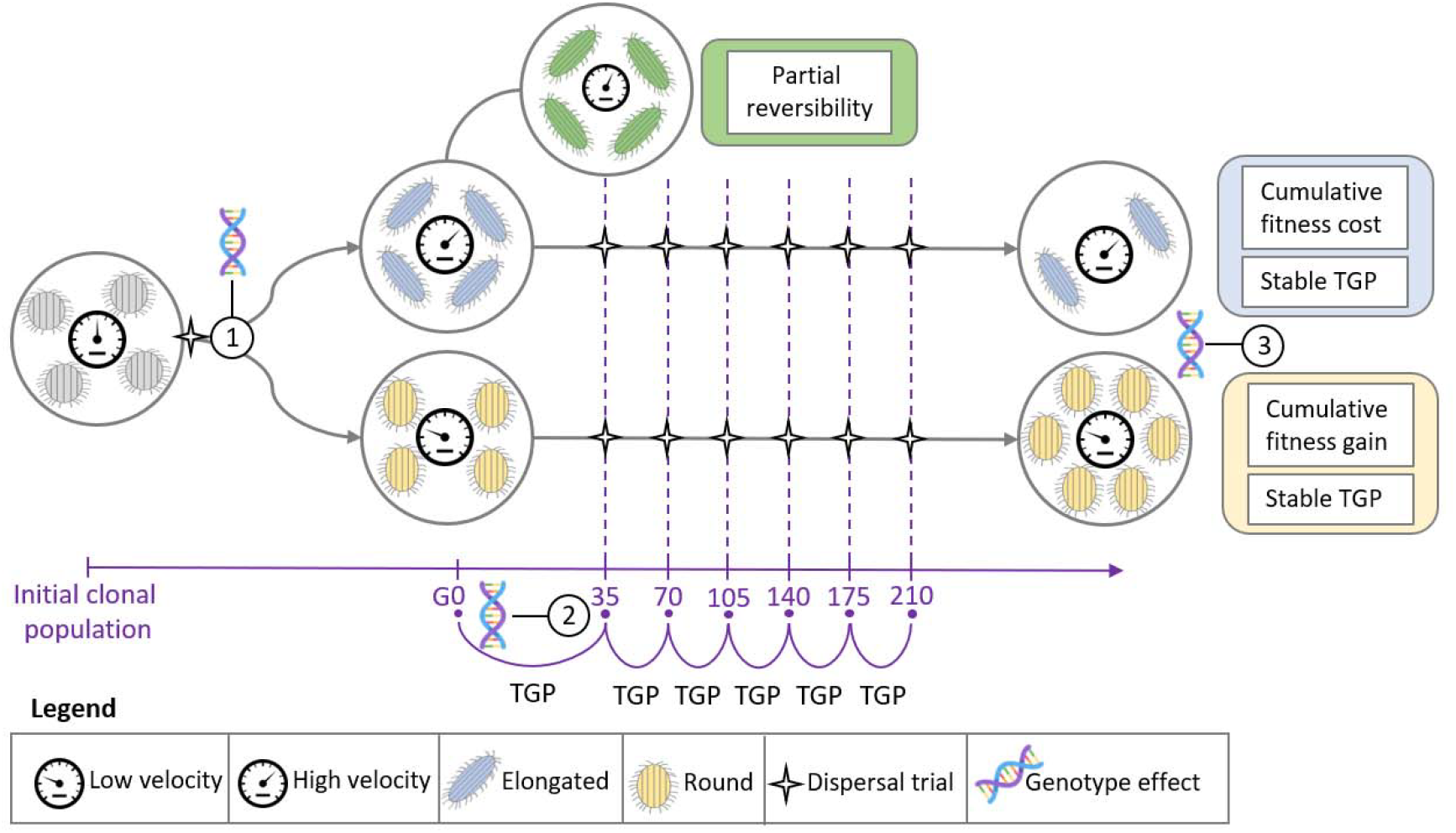
Transgenerational plasticity for dispersal and its cost in Tetrahymena thermophila. At generation 0 (G0), the initial dispersal trial is performed (dispersal trials are represented by the black stars). After the first trial, cells are more elongated and swim faster than in mother cultures, but dispersing cells (in blue) have a more elongated shape and a higher velocity than non-dispersing cells (yellow) due to plastic changes within genotypes. The strength of phenotypic differences between dispersing and non-dispersing cells differ between genotypes (1). The dispersal status of the ancestor affects the phenotype of descendants: cells with a dispersing ancestor conserve a dispersing-like phenotype (elongated and fast) via transgenerational plasticity during whole the experiment. Yet, the strength of this effect depends on cell genotype (2). These phenotypic changes are only partially reversible (in green) after ~35 generations in common garden (velocity slightly decreases while elongation slightly increases, fitness is stable). The number of dispersal trials experienced by the ancestors of a cell does not affect its phenotype: the effect of transgenerational plasticity is not gradual. Indeed, the phenotypic switches appear at the first trial and are then maintained throughout the experiment. By contrast, cells with dispersing ancestors experience a gradual decrease in fitness along with the number of dispersal trials experienced by their ancestors. Likewise, the fitness of cells with non-dispersing ancestors increases with the number of dispersal trials experienced. Genotype modulates this fitness effects of transgenerational plasticity for dispersal (3).

We also observed a cumulative fitness cost associated with dispersal, while non-dispersing cells increased their fitness. To the best of our knowledge, cumulative fitness costs of plasticity across ~200 generations have never been described in the context of dispersal. While fitness dynamics should be built on more time points and for more generations in the future, our result is of utmost importance because differential costs and benefits associated with dispersal strategies can drive their coexistence (Bonte et al. 2012). *T. thermophila* thus offers an interesting system to test a series of predictions and calibrate models on the role of plasticity, dispersal, and their costs and benefits on eco-evolutionary dynamics (*e.g.*, Scheiner et al. 2012, Scheiner et al. 2017). Future work in ciliates and other taxa should also determine the tipping points at which TGP costs of dispersal would alter colonisation and/or (meta)population dynamics (Doebeli & Ruxton 1997).

At first glance, trait variance explained by our ancestor dispersal status might appear low (from 1 to 13% depending on the trait and genotype). However, dispersal is a multifaceted process for which tens (or more) phenotypic traits are involved (Clobert et al. 2009). Therefore, it might not be surprising that, working on only four candidate traits, we measured moderate responses in our simple experimental conditions. Besides, cell shape and velocity are involved in numerous other fundamental cell functions (*e.g.*, feeding, mating, osmoregulation), which certainly impose constraints on their variance. Finally, fitness differed in mean by 9% between dispersing and non-dispersing cells, suggesting that transgenerational dispersal plasticity can strongly impact evolutionary dynamics.

In our experiment, plastic changes were only partially reversible between the dispersal trials. Velocity measured just after each trial was weakly lower after ~35 generations in common garden, but still higher in dispersing cells with a dispersing ancestor than in dispersing cells with a non-dispersing ancestor. Dispersing cells with a dispersing ancestor were even more elongated after the common garden, which might be due to the dispersal treatment itself, or to phenotypic differences potentially observed between growth stages (Taylor et al. 1976). Finally, the fitness difference between dispersing and non-dispersing cells was not affected by the common garden. Such limited reversibility of phenotypes suggests either that the mechanisms responsible for this dispersal plasticity present a time-lag to fully reverse the phenotypes, or that the environmental cues triggering the phenotypic reversibility are not entirely reliable (the two hypotheses being non-exclusive).

In absence of substantial genetic variation within the cell lines, the described inheritance of dispersal-related traits should rely on non-genetic factors causing transgenerational modifications of gene expression (Devanapally et al. 2015). In *T. thermophila*, epigenetic mechanisms as DNA methylation (Chung & Yao 2012), microRNA (Mochizuki 2012), or histone modifications (Morris et al. 2007) might allow the transmission of changes in cell shape and velocity across clonal generations. As the ciliate somatic genome is highly polyploidized (~45 copies in *T. thermophila*, Doerder et al. 1992), epigenetic modifications induced before or during the dispersal process could cause differences in the expression of specific copies of homeologous genes coding for dispersal-related traits (Liu & Adams 2007). In our experimental design, the absence of sexual reproduction, and therefore the lack of meiotic reprogramming of epimarks, should facilitate the transgenerational inheritance of epigenetic variants regulating the expression of homeologous genes (Heard & Martienssen 2014), and should thus foster the TGP for dispersal. In *T. thermophila*, copy number variation can generate adaptive plastic responses under stressful conditions with a time lag of at least a few generations (di Fransisco et al. 2018). While it should be excluded that copy number variation explains the initial phenotypic changes in our experiment (cells are different from mother cultures in both dispersing and non-dispersing lines at the first trial), it is possible that epigenetic modifications followed by copy number variations act in concert to maintain the observed TGP. The time lag associated with copy number regulation could then account for the partial reversibility of phenotypes observed, as well as progressive elimination of mRNA, microRNA, or other intracellular molecules potentially responsible for TGP through cell divisions.

Our study showed that genetic background explained the differential persistence of dispersal phenotypes during ~35 asexual generations (Fig.2). As well, cell genotype significantly modulated the transgenerational fitness consequences, where the more canalized genotypes for dispersal (*i.e.* those presenting the lowest plastic response, D3 and D6) experiencing more costs when regularly confronted to dispersal trials. This suggests that genotypes able to plastically express specialized dispersing phenotypes have evolved mechanisms to reduce the associated costs. Our results therefore revealed that G × E interactions drive the TGP for dispersal and its cost in *T. thermophila*. Phenotypic trade-offs are usually observed in the context of dispersal (Bonte et al. 2012), but we highlight here an original dependency on the genetic background. A genetic control of TGP has rarely been observed (see however Devanapally et al. 2015, Vu et al. 2015), and could be caused by the genetic determinism of epimarks’ transgenerational inheritance. Indeed, methylation variation are usually strongly associated with genetic variants in both *cis* and *trans* (Dubin et al. 2015, Zaghlool et al. 2016), facilitating or constraining the transmission of epimarks over generations (Richards 2006). In the future, comparisons between epigenomes and transcriptomes of the tested genotypes should provide mechanistic answers. It should also be helpful to understand if the parallelism found between the biological replicates of each genotype and for some traits between genotypes (models all include replicates and genotypes as variables) relies on similar molecular mechanisms.

To conclude, our study provides a first evidence of the role of genetic background in the TPG and associated cost in a dispersal context. It emphasizes the tremendous importance of G × E interactions in the ability of organisms to transmit phenotypic variations induced by the environment across generations, shedding light on the importance of intraspecific genetic variation in ecological and evolutionary dynamics (*e.g.*, Raffard et al. 2018). Our results outline that genotype-dependent TGP likely plays a critical role in the evolution of dispersal, a major eco-evolutionary force that determines the migration-drift and migration-selection equilibria in natural populations (Slatkin 1987, Lenormand 2002). Genetically-controlled TGP for dispersal could also be a central mechanism in biological invasions by allowing a rapid phenotypic specialization maximizing colonization success and speed (Perkins et al. 2013, Ochocki & Miller 2017), despite a low genetic polymorphism caused by serial founder effects (Excoffier et al. 2009). More broadly, genotype-dependent TGP could facilitate a rapid adjustment to sudden environmental changes, such as climate change, especially when standing genetic variation is low and the chances of beneficial mutation recruitment are small. In this regard, it might be of high concern to determine if the degree of parallelism measured here can also be observed at the inter-specific level. This would help quantify the importance of plastic mechanisms in biodiversity response to environmental changes.

## Material and methods

### Model species and culture conditions

*Tetrahymena thermophila* is a 30-to 50-µm ciliated unicellular eukaryote naturally living in freshwater ponds in North America, which alternates sexual and asexual phases depending on environmental conditions. The species is a model organism in cell and molecular biology, and its maintenance under laboratory conditions benefits from decades of experience (Collins 2012). We used four genotypes originally sampled and kindly provided by F. P. Doerder between 2002 and 2008 in North America (genotype D3, D4, D6, and D9; Pennekamp et al. 2014), and bred uniquely under clonal conditions. Before and during the experiment, cells were all cultivated in the same standard conditions: 23°C in climatic chambers in 0.3X synthetic liquid growth media (0.6% Difco proteose peptone, 0.6% yeast extract) as described in previous studies (Fjerdingstad et al. 2007, Schtickzelle et al. 2009, Jacob et al. 2015). In these conditions, the cell division time is around 4-6 hours (~5 generations per day). All manipulations were performed in sterile conditions under a laminar flow hood.

### Protocol of successive dispersal trials

We performed an experimental procedure of repeated dispersal trials to investigate how phenotype of cells is affected by the dispersal status of their ancestors and how the number of experienced trials affects the phenotype of descendants. Dispersal trials were performed using standard connected microcosms composed of two habitat patches consisting of 1.5 ml microtubes connected by a corridor made of 4 mm internal diameter, 2.5-cm long silicone tube (Jacob et al. 2016). These laboratory conditions proved useful to study many aspects of dispersal such as, *e.g.*, the architecture of dispersal syndromes, the causes of dispersal (Pennekamp et al. 2014, Fronhofer et al. 2018), the cooperation-colonization trade-off (Jacob et al. 2016), range expansions (Fronhofer & Altermatt 2015, Fronhofer et al. 2017), or (meta)population and community dynamics (Fox et al. 2014, Jacob et al. 2019). For a dispersal trial, a fraction of ~100,000 cells were placed in one of the two patches, called the departure patch while corridors were closed with clamps. Then, corridors were opened and cells were therefore allowed to either stay in the departure patch or disperse to the other patch, called arrival patch, over a 4-hours period. After this period, the corridors were clamped and samples from the two populations of cells (dispersing in the arrival patch and non-dispersing in the departure patch) were pipetted to inoculate a new separately growing population.

For the four genotypes, we isolated by hand-pipetting one mother cell that reproduced clonally over a 7-day period in one 2 ml well of a 24-well plate. From this initial mother-culture, we made five replicates (*i.e*. initial populations) that were cultivated over another 7-days period (~35 cell divisions; see **Supplementary material**, **Fig.S1**). Then, these 20 populations (*i.e*., five replicates in four genotypes) experienced an initial dispersal trial (*tr*0) that allowed producing one subpopulation with dispersing ancestors and one subpopulation with non-dispersing ancestors. The two subpopulations were subjected to a new dispersal trial every seven days to obtain a total of six trials (*tr*1 to *tr*6). Over the successive trials, we serially kept and cultivated dispersing cells and non-dispersing cells in the subpopulations with dispersing and non-dispersing ancestors respectively (**Supplementary material, Fig. S1**).

### Phenotype and fitness measurements

The four phenotypic traits (morphology: cell size and shape; movement: velocity and linearity) were measured in initial populations. Then, from trials *tr*0 to *tr*6, the same traits were measured just ‘before’ and ‘after’ each dispersal trial. The ‘before’ measurement was used to quantify traits after 7 days, *i.e.*, around 35 generations, in common garden conditions (standard medium without dispersal possibility). The ‘after’ measurement was used to quantify traits at the exact time of dispersal. Cell size (area in µm²) and shape (cell major/minor axis ratio of a fitted ellipse), as well as velocity (µm/s) and movement linearity (distance in straight line/effective distance covered), were measured using on automated analysis of digital images and videos (Pennekamp & Schtickzelle 2013, Pennekamp et al. 2015). For each sample of cells, we considered five technical replicates (10 μl) pipetted into one chamber of a multi-chambered counting slide (Kima precision cell 301890), and took digital pictures under dark-field microscopy (Pennekamp & Schtickzelle 2013). Data from the five technical replicates were pooled in all analyses. We used ImageJ (version 1.47, National Institutes of Health, USA) and BEMOVI (Pennekamp et al. 2015) softwares to measure morphological and movement variables. Using the same program, we calculated dispersal rates at each dispersal trial by quantifying cell density in the patches of departure and arrival using the automated analysis of digital images described above (see Pennekamp et al. 2014).

We measured cell fitness in initial populations and after the dispersal trial (for both dispersing and non-dispersing cells at three dispersal trials (*tr*0, *tr*1, and *tr*6) using standard population growth analyses. Small numbers of cells (~100 cells) were transferred in four technical replicates into 96-well plates filled with 250 µl of fresh growth media. Cultures were maintained at 23 °C and absorbance measurements at 550 nm were performed every 2 h for 2 weeks using an automated microplate reader (Tecan Infinite Spectrophotometer with a Connect robotized arm). We then computed the growth rate as the maximum slope of population growth through time and the maximal population density as the density reached at the plateau by smoothing the absorbance data using general additive model (gam package; Hastie 2018), and fitting a spline-based growth curve using the *grofit* package of R (*gcfit* function; Kahm et al. 2010). For simplicity, we present results on growth rate, the most frequently used fitness proxy (Orr 2009), given that results were qualitatively similar using maximal density (data not shown).

### Statistical analyses

#### Trait covariation (models 1 of Table S1)

First, we assessed the initial relationships between the four phenotypic traits (i.e., cell shape, cell size, movement velocity and linearity). We used phenotypic measurements recorded prior the first dispersal trial (*tr*0) to assess between-traits covariation pattern. A linear mixed model was used to examine the correlation between each pair of traits. One of the two phenotypic trait was treated as the dependent variable whereas the other was introduced in the model as an explanatory term. The dependent variable was log-transformed and the explanatory variable was z-scored. The strain and the replicate were introduced as random effects in the model. For all analyses implicating linear mixed models, we used restricted maximum likelihood optimization. Normality of the residuals was examined graphically using a quantile–quantile plot. We used a likelihood ratio test to assess the significance of the relationship, i.e. comparing the models with and without the explanatory term. We calculated marginal *R*^2^ to quantify the proportion of variation explained by the explanatory variable only.

#### Effect of genotype on cell phenotype and fitness (models 2 of Table S1)

We evaluated the influence of the cell genetic background on the four phenotypic traits and cell growth rate (i.e., a proxy of cell fitness) before the first dispersal trial at *tr*0. We used linear mixed models in which the log-transformed phenotypic traits were introduced as dependent variables, cell genotype as the explanatory variable (i.e. a discrete variable with four modalities) and the replicates as random effects. We used a similar procedure to examine the effect of genotype on fitness at *tr*0.

#### Dispersal syndrome and dispersal-related fitness cost (models 3 of Table S1)

We examined how morphology and movement behavior correlate with cell dispersal status after the first dispersal trial (*tr*0). We made general analysis where all genotypes were combined. We used linear mixed models where the log-transformed phenotypic traits were introduced as dependent variables, the cell dispersal status as the explanatory variable (*i.e.*, a discrete variable with two modalities, dispersing *vs* non-dispersing), and the genotype and replicate as random effects.

#### Effect of genotype and ancestor dispersal status on descendant phenotype and fitness (models 1 of Table S2)

We examined the effect of ancestor dispersal status (dispersing vs non-dispersing lines coded as a discrete variable) on phenotypic trait and fitness of descendants after ~ 35 cell divisions (*i.e.*, 7 days). First, we made a general analysis using linear mixed models where log-transformed phenotypic traits and fitness were introduced as dependent variables, and the ancestor dispersal status as discrete explanatory variable. The genotype, the replicate, and the number of dispersal trials experienced by ancestor were included in the model as random effects (**models 1.1** of **Table S2**). Then, we conducted a partial analysis where we analyzed the four genotypes separately (**models 1.2** of **Table S2**).

#### Effect of the number of successive dispersal trials on descendant phenotype and fitness (models 2 of Table S2)

We retrieved the same linear mixed models used to investigate the effect of ancestor dispersal status on phenotype and fitness, but the number of dispersal trials experienced was removed from the random effects and introduced in the fixed part of the model. For the phenotypic traits, the number of dispersal trials (from 0 to 6) was incorporated as a continuous variable. We tested additive and interactive effects (ancestor dispersal status × number trials) of the variable, and considered both linear and logarithmic relationships; a likelihood ratio test has been performed to compare the two relationships. For cell fitness, the number of dispersal trials (0, 1, and 6) was entered in the model as discrete variable, and both additive and interactive effects were examined.

### Reversibility of transgenerational plastic changes (models 3 of Table S2)

We examined how stable were the transgenerational changes of cell phenotype by comparing the phenotype of cells measured after each dispersal trial and the phenotype of their descendants after ~ 35 asexual generations in common garden (**model 3 of Table S2**) in dispersing lines. We used linear mixed models where log-transformed phenotypic traits and fitness were introduced as dependent variables, and the type of cell as explanatory variable. We included the genotype, the replicate, and the number of dispersal trials experienced by ancestor as random effects.

## Acknowledgements

H.C. is supported by a Vanier-Banting postdoctoral fellowship. This work was supported by the French Laboratory of Excellence project TULIP (ANR-10-LABX-41). The authors warmly thank Julien Cote and Kyle Wellband for helpful discussion on this manuscript.

