## Supplementary_material for "Transgenerational dispersal plasticity and its fitness consequences are under genetic control"

**Genetic background and transgenerational plasticity drive dispersal evolution in a protist**

**Supplementary material**

**Table S1**. Linear models and linear mixed models used for the analyses focusing on the observations made at *tr*0. The level of observation (cell or population), the sample size (number of observations) are presented. We also present for each model the response variable, the random effects, and the fixed effects. We used observations made at cell level (except for population growth rate) to increase our statistical power – the number of observations at the population level was too limited to run the analyses.

| Observation level | Sample size | Response variable | Fixed effects | Random effects |
| --- | --- | --- | --- | --- |
| 1. Phenotypic trait covariation | | | | |
| Cell | 5611 | shape | size | genotype + replicate |
| Cell | 5611 | shape | velocity | genotype + replicate |
| Cell | 5611 | velocity | linearity | genotype + replicate |
| Cell | 5611 | velocity | size | genotype + replicate |
| Cell | 5611 | linearity | size | genotype + replicate |
| Population | 20 | growth rate | shape | genotype + technical replicate |
| Population | 20 | growth rate | velocity | genotype + technical replicate |
| Population | 20 | growth rate | size | genotype + technical replicate |
| Population | 20 | growth rate | linearity | genotype + technical replicate |
| 2. Effect of genotype on cell phenotype and fitness | | | | |
| Cell | 5611 | shape | genotype | replicate |
| Cell | 5611 | size | genotype | replicate |
| Cell | 5611 | velocity | genotype | replicate |
| Cell | 5611 | linearity | genotype | replicate |
| Population | 20 | growth rate | genotype | technical replicate |
| 3. Dispersal syndrome and dispersal-related fitness cost | | | | |
| Cell | 5611 | shape | cell dispersal status | genotype + replicate |
| Cell | 5611 | size | cell dispersal status | genotype + replicate |
| Cell | 5611 | velocity | cell dispersal status | genotype + replicate |
| Cell | 5611 | linearity | cell dispersal status | genotype |
| Population | 160 | growth rate | cell dispersal status | genotype + replicate |

**Table S2**. Linear models and linear mixed models used for the analyses focusing on the observations made at all dispersal trials (from *tr*0 to *tr*6). We used observations made at the population level (i.e., the mean of phenotypic trait recorded in the cell population) to avoid the statistical issues associated with large sample size (> 350,000 observations for the analyses of step 2 and 3). We also present for each model the response variable, the random effects, and the fixed effects.

| Observation level | Sample size | Response variable | Fixed effects | Random effects |
| --- | --- | --- | --- | --- |
| 1. Ancestor dispersal status on descendant phenotype and fitness (general time-constant analyses) | | | | |
| 1.1. General analysis (all genotypes combined) | | | | |
| Population | 240 | shape | ancestor dispersal status | genotype + replicate + time |
| Population | 240 | velocity | ancestor dispersal status | genotype + replicate + time |
| Population | 660 | growth rate | ancestor dispersal status | genotype + replicate + technical replicate + time |
| 1.2. Partial analysis (per genotype) | | | | |
| Population | genotype-dependent | shape | ancestor dispersal status | replicate + time |
| Population | genotype-dependent | velocity | ancestor dispersal status | replicate + time |
| Population | genotype-dependent | growth rate | ancestor dispersal status | replicate + technical replicate + time |
| 2. Ancestor dispersal status on descendant phenotype and fitness (general time-specific analyses) | | | | |
| Population | 280 (*tr*0 added) | shape | ancestor dispersal status × time | genotype + replicate |
| Population | 280 (*tr*0 added) | velocity | ancestor dispersal status × time | genotype + replicate |
| Population | 660 (*tr*0 added) | growth rate | ancestor dispersal status × time | genotype + replicate + technical replicate |
| 3. Reversibility of transgenerational plastic changes | | | | |
| Population | 240 | shape | type of cells (ancestor *vs* descendants) | genotype + replicate + time |
| Population | 240 | velocity | type of cells (ancestor *vs* descendants) | genotype + replicate + time |
| Population | 400 | growth rate | type of cells (ancestor *vs* descendants) | genotype + replicate + time |

**Table S3**. Correlation between phenotypic traits in mother cultures just before trial *tr*0.

| Trait association | $R^{2}$ | $\chi^{2}$ | P-value |
| --- | --- | --- | --- |
| velocity – size | < 0.0001 | 10.88 | 0.0001 |
| velocity – shape | 0.17 | 71259.00 | < 0.0001 |
| linearity – size | 0.01 | 2757.50 | < 0.0001 |
| linearity – shape | 0.07 | 18588.00 | < 0.0001 |
| velocity – linearity | 0.09 | 43787.00 | < 0.0001 |
| size – shape | 0.01 | 10445.00 | < 0.0001 |

**Table S4**. Effect of the interaction between ancestor dispersal status and number of selection events on phenotypic traits.

| Trait association | $\chi^{2}$ | P-value |
| --- | --- | --- |
| VELOCITY |  |  |
| Status × number of events | 2.13 | 0.14 |
| SHAPE |  |  |
| Status × number of events | 1.63 | 0.20 |

**Table S5**. Effect of ancestor dispersal status on velocity and shape in the four studied genotypes.

| Trait association | $R^{2}$ | $\chi^{2}$ | P-value |
| --- | --- | --- | --- |
| VELOCITY |  |  |  |
| D3 | 0.10 | 2.15 | 0.14 |
| D4 | 0.06 | 3.96 | 0.04 |
| D6 | 0.003 | 0.72 | 0.77 |
| D9 | 0.13 | 6.53 | 0.01 |
| SHAPE |  |  |  |
| D3 | 0.01 | 1.56 | 0.45 |
| D4 | 0.01 | 1.66 | 0.21 |
| D6 | 0.001 | 0.03 | 0.88 |
| D9 | 0.11 | 8.13 | 0.04 |

**Table S62**. Effect of ancestor dispersal status on growth rate.

| Trait association | $R^{2}$ | $\chi^{2}$ | P-value |
| --- | --- | --- | --- |
| D3 | 0.13 | 54.04 | <0.0001 |
| D4 | 0.10 | 27.71 | <0.0001 |
| D6 | 0.06 | 133.01 | <0.0001 |
| D9 | 0.05 | 73.84 | <0.0001 |

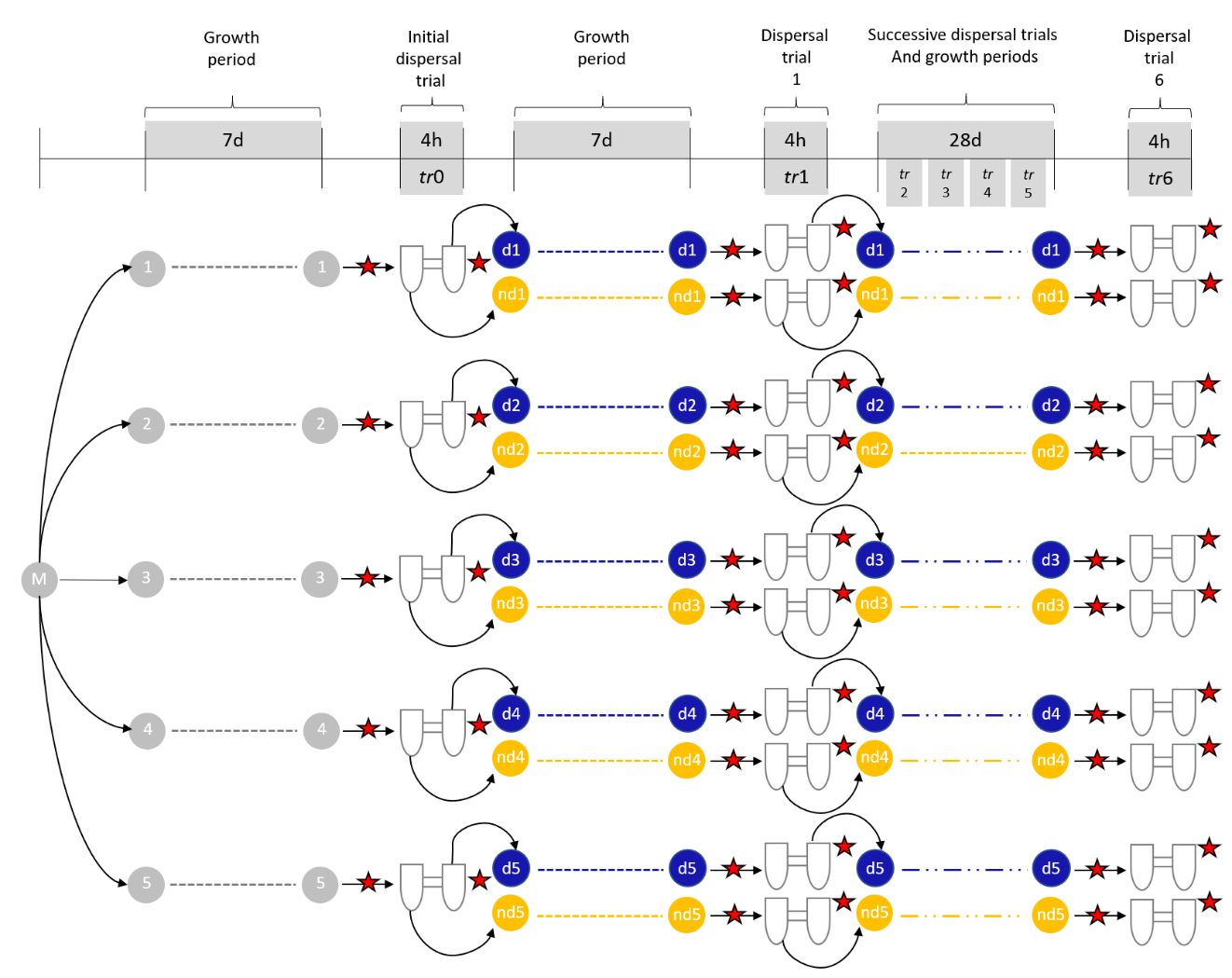

**Fig.S1**. Design of artificial selection. For the four cell genotypes (D3, D4, D6, and D9), we generated five initial replicated populations (1 to 5) from a mother culture (M) obtained from the isolation of a single cell (*i.e.,* isogenic population). For simplicity, only one genotype is represented. The five replicated initial populations were cultivated over a 7-days period. Next, we performed the initial dispersal trial (*tr*0) from which we produced one subpopulation (from 1 to 5) with dispersing ancestors (*d*) and one subpopulation with non-dispersing ancestors (*nd*). After a 7-days period of growth, we performed another dispersal trial (*tr*1). We kept and cultivated dispersing cells and non-dispersing cells for *d* and *nd* subpopulations respectively. This procedure was repeated five more times to obtain a total of six dispersal trials. Phenotypic measures (cell size, cell shape, movement linearity and velocity) and fitness measures (growth rate) were performed just before and just after dispersal trials (red stars).

**Phenotypic changes before and after the initial trial**

We investigated whether the shape and velocity of descendants differed from the phenotype of their ancestor prior to the initial dispersal trial (~ 35 generations before). We performed separated analyses using individual data (45,512 observations). We built models where phenotypic traits measured prior the initial (*tr*0) and the subsequent (*tr*1) dispersal trials were included as the response variable. The time (*tr*0 vs *tr*1) and the ancestor dispersal status (D and ND) were merged into a discrete covariate with three modalities: cells prior initial dispersal trial *tr*0, cells prior dispersal trial *tr*1 with a non-dispersing ancestor at *tr*0 (ND), and cells prior dispersal trial *tr*1 with a dispersing ancestor at *tr*0 (D). Genotype identity and replicates were included in the model as random effects.

Our analyses showed that cells at *tr*1 had a longer shape than cells prior *tr*0 (I as the intercept, 1.01±0.04; D, slope coefficient: 0.07±0.02; ND, slope coefficient: 0.05±0.02) regardless of the phenotype of their ancestor at the dispersal trial *tr*0 (LR test: $R^{2}$ = 0.04, $\chi^{2}$ = 110.00, *p* < 0.0001). As shown in the analyses of step 1 and 2 (see Table S2), they also indicate that cells with a dispersing ancestor had longer shape than cells with a non-dispersing ancestor (see the slope coefficients presented above).

Our analyses also revealed that cells at *tr*1 had a higher velocity than cells prior *tr*0 (I as the intercept, 5.43±0.12; D, slope coefficient: 0.43±0.11; ND, slope coefficient: 0.22±0.12) regardless of the phenotype of their ancestor (D or ND) at the dispersal trial *tr*0 (LR test: $R^{2}$ = 0.10, $\chi^{2}$ = 2513.00, *p* < 0.0001). As demonstrated in the analyses of step 1 and 2 (see Table S2), they also showed that cells with a dispersing ancestor had higher velocity than cells with a non-dispersing ancestor.

To summarise, those additional results indicate that cells with dispersing and non-dispersing ancestor have a longer shape and a higher velocity than cells prior the initial dispersal trial. The difference in trait variance between mother cultures and tested cells rules out the possibility that the phenotypic differences between dispersing and non-dispersing cells results from the spatial segregation of cells with residual genetic variation within populations. Indeed, while we have initiated cultures with a single mother cells, the somatic genome of *T. thermophila* is ~45 polyploidized, with random segregation of chromosome copies at each division. It might then exist low residual genetic variability in mother cultures. However, in this case, the variance in traits in mother cultures and in the pooled dispersing and non-dispersing cells should be similar, which is not the case. At the methodological level, our results also indicate that performing the analyses with individual (the present analyses) or population data (analyses steps 1 and 2, Table S2) provides relatively similar results and conclusions concerning the effect of ancestor dispersal status on descendant phenotype.
